# Somatic cell nuclear transfer in non-enucleated goldfish oocytes: understanding DNA fate during meiosis resumption and first cellular division

**DOI:** 10.1101/630194

**Authors:** Charlène Rouillon, Alexandra Depincé, Nathalie Chênais, Pierre-Yves Le Bail, Catherine Labbé

## Abstract

Nuclear transfer consists in injecting a somatic nucleus carrying valuable genetic information into a recipient oocyte to sire a diploid offspring who bears the genome of interest. It requires that the oocyte (maternal) DNA is removed. In fish, because enucleation is difficult to achieve, non-enucleated oocytes are often used and disappearance of the maternal DNA was reported in some clones. The present work explore which cellular events explain spontaneous erasure of maternal DNA, as mastering this phenomenon would circumvent the painstaking procedure of fish oocyte enucleation. The fate of the somatic and maternal DNA during meiosis resumption and first cell cycle was studied using DNA labeling and immunofluorescence in goldfish clones. Maternal DNA was always found as an intact metaphase within the oocyte, and polar body extrusion was minimally affected after meiosis resumption. During the first cell cycle, only 40 % of the clones displayed symmetric cleavage, and these symmetric clones contributed to 80 % of those surviving at hatching. Maternal DNA was often fragmented and located under the cleavage furrow. The somatic DNA was organized either into a normal mitotic spindle or abnormal multinuclear spindle. Scenarios matching the DNA behavior and the embryo fate are proposed.

## Introduction

Somatic cell nuclear transfer (SCNT) consists in injecting a donor somatic nucleus into a recipient oocyte to obtain a clone that carries the genome of the donor animal^1–3^.This technology allows the restoration of valuable genetic resources from somatic material when both sperm and oocytes, or embryos, are unavailable. This is the case in fish, where the most recognized support for the preservation of genetic resources is solely the cryopreserved sperm^4^, but where neither oocytes nor embryos can be cryopreserved^5^. In this context, cryopreserved fin cells, which carry the genome of both parents, are particularly beneficial for regenerating a breeder^6^. Somatic diploid cells are easily collected from fins by non-invasive methods regardless the characteristics of the fish (age, maturity stage, sex, conservation status). Somatic cells are easy to culture *in vitro* and to cryopreserve even under difficult experimental conditions^7,8^.

After SCNT, the production of offsprings based on the genome of the donor only requires that the contribution of the oocyte (maternal) genome is prevented. In mammals, this is achieved by enucleating the recipient oocyte before performing the SCNT. In a majority of fish, the oocyte structure makes enucleation very difficult: oocytes are large and opaque, they contain bulky nutritional reserves (the yolk), a dense cytoplasm (the ooplasm), and a thick protective envelope around the oocyte (the chorion)^9,10^. These characteristics prevent the vizualization of maternal genome by transparency and its aspiration for enucleation. Most authors overcome these difficulties by activating oocytes and taking the second polar body as a guide for putative localization of maternal pronucleus^11–13^. However, these oocytes are less suitable for donor DNA reprogramming than non-activated oocytes^14^, and aspiration of the female pronucleus is associated with loss of essential developmental factors such as maternal mRNAs, mitochondria and proteins. Thanks to highly focused laser irradiation of the non-activated oocyte, the Cibelli group^15,16^ succeeded in inactivating the maternal metaphase at a more appropriate recipient stage, but adaptation of this method to species other than zebrafish was never reported, likely because of the difficult tuning of the laser on different egg types. Overall, in addition to be time consuming, fish enucleation is a very problematic issue for the success of nuclear transfer and embryonic development of the clone. For this reason, several authors have attempted to carry out nuclear transfer without any inactivation or elimination of maternal DNA. It is interesting that in goldfish, zebrafish, weatherfish and medaka, such a protocole avoiding the non-enucleation step still allows the development of clones carrying only the genome of the donor^17–22^.

Understanding the mechanisms responsible for the spontaneous loss of oocyte DNA and/or the possible interference induced by retained maternal DNA during early development would help to promote the development of clones from donor origin only. However, this issue is hampered in fish by the lack of knowledge about cellular events that occur after SCNT. It is not known for example whether meiosis resumes normally, and how the clone ploidy is established. Ovulated oocytes bear a condensed maternal DNA maintained in a metaphase plate (MII stage) up to fertilization, when meiosis resumption triggers the second polar body extrusion and maternal genome haploidization^10^. Polar body extrusion has never been studied in fish after SCNT, and although few studies explored clone ploidy^18–20,23^, the remodelling patterns of the maternal and somatic chromatins in the clones during the first cell cycle are not known.

In this context, the objective of our study was to understand the fate of maternal DNA and the mechanism of spontaneous enucleation in clones, and to characterize the interplay between somatic and maternal DNA. First, we explored the organization of maternal and somatic DNA after donor cell injection into the MII stage oocyte, in order to evaluate the disturbance profile and location of DNA from both origins. After oocyte activation, we analyzed the extrusion pattern of the polar body to identify how DNAs were handled by the oocyte environment during meiosis resumption. Then, we characterized the organization of the blastomere during the first cell division and identified the fate of DNA from both origins at this stage. Finally, we discussed how these mitotic figures can explain the spontaneous neutralization of maternal DNA as well as the many embryonic defects observed in clones.

## Results

### Maternal and somatic DNA fates after injection into the MII oocyte

Characterization of the various compartments in control oocytes was necessary in order to identify and localize the maternal and somatic DNA in the clones. In goldfish, the opacity of the chorion and light refraction of the oocytes prevented any live analysis of the whole oocytes. Instead, all samples had to be fixed and analyzed on histological sections. Oocyte structure (Fig. 1.A-B) was very consistent: the chorion was observed as a thick membrane with a crenellated shape. We hypothesize that this shape would permit expansion of the chorion upon activation, to allow the formation of the perivitelline space. The micropyle (sperm entry point) was always found as a canal-like invagination of the chorion. Its depth depended on the section angle, but it always penetrated a homogeneous area that was assigned to the ooplasm of the animal pole. Under the chorion and all around the oocyte, the cortical granules were embedded in a thin cytoplasmic layer. They were distinguished from yolk droplets by their transparency under UV light, while yolk droplets appeared as transparent globules under visible light (Fig. 1.A,C). The density of oocyte yolk droplets increased as the sections deepened towards the vegetative pole. Interestingly, in 100% of control oocytes (n=18), maternal DNA was not found at the bottom of the micropyle, but against the side of the micropylar canal, in the ooplasm, and it was always organized in a metaphase plate (Fig. 1.C).

**Figure 1.**
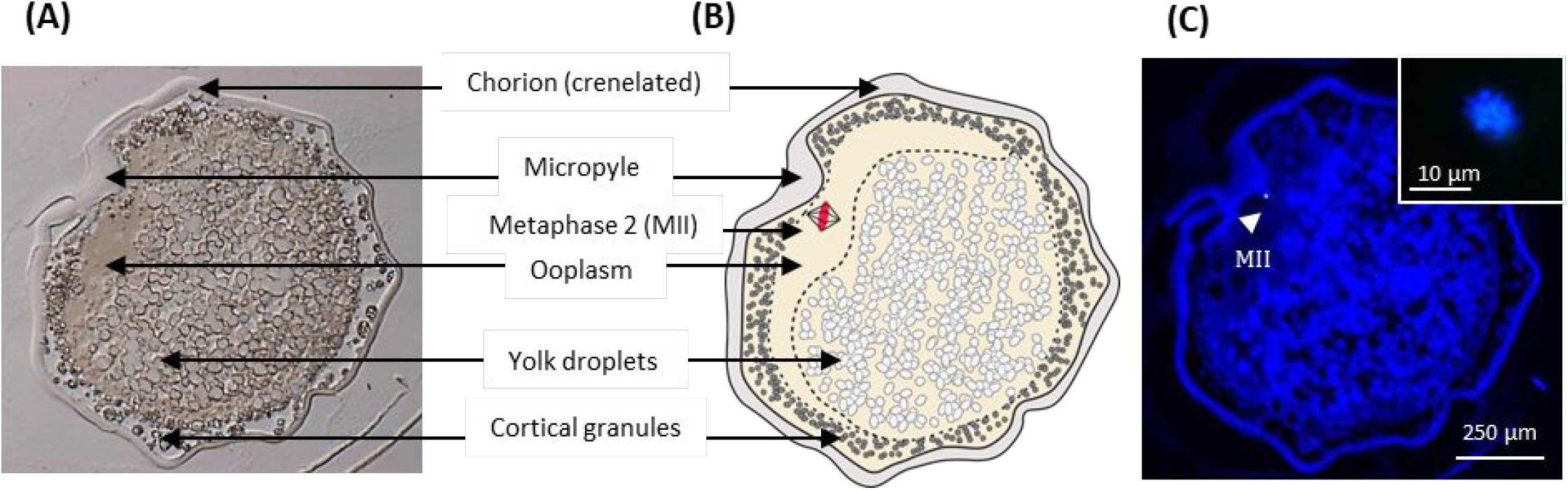
Structure of a non-activated control oocyte and location of the maternal DNA. Control oocytes were fixed, cut into 7 μm sections and DNA was stained with Hoechst 33342. (A) Image of a section of a non-activated oocyte observed under visible light. (B) Schematic representation of the oocyte compartments, validated by the observation of successive sections from 18 oocytes. (C) Image of the oocyte observed under UV-light, arrowhead shows the maternal DNA condensed in a metaphase plate (MII). The MII was located against the micropylar canal. Inset represents the MII observed in confocal microscopy.

In the clones, maternal DNA was present in every oocytes (100%, n=28 clones), and it was located against the micropylar canal (Fig. 2.A,E), like in the control oocytes. This DNA was always condensed in a metaphase plate (MII stage) that was not different from the one of non-activated control oocytes. In addition, even if the oocyte membrane was perforated during injection through the micropylar canal, the entire structure of the oocyte was similar to the one of the non-activated control oocytes.

**Figure 2.**
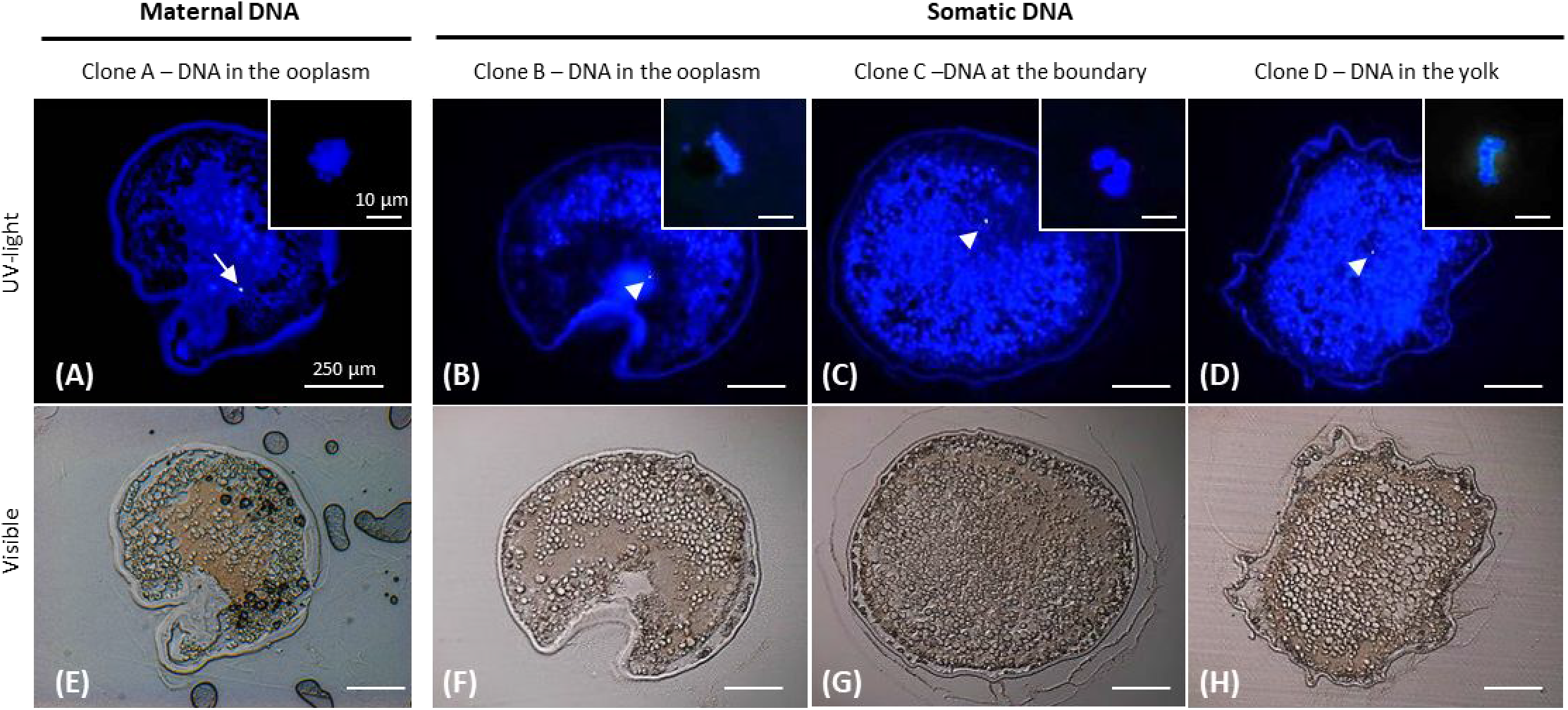
Maternal and Somatic DNA location into non-activated clones. Non-activated clones after somatic DNA injection were fixed, cut into 7 μm sections and DNA was stained with Hoechst 33342. Images of a section of a clone (Clone A) observed (A) under UV- and (E) visible light. The arrow shows the maternal DNA in a metaphase plate (MII), located against the micropylar canal, as in control oocytes. This example (clone A) is representative of 28 clones. Images of sections of 3 different clones (Clones B to D) observed under (B-D) UV-light or (F-H) visible light, showing the various locations of the somatic DNA : (B,F) into the ooplasm, (C,G) at the yolk border, and (D,H) into the yolk reserve. Those three clones are representative of the 16 clones in which the somatic DNA was observed. Arrowheads show the somatic DNA. Insets represent the maternal or somatic DNA observed by confocal microscopy.

Such a consistency in location and shape of the maternal DNA allowed us to precisely distinguish it from the somatic DNA. Somatic DNA was observed in 57% of non-activated clones (n=16/28). Although care was taken to inject the somatic cell just under the plasma membrane at the bottom of the micropylar canal, its DNA was found in different compartments within the oocytes (Fig. 2): most of the somatic DNA was found in the ooplasm (37.5%) (Fig. 2.B,F), or at the boundary between the ooplasm and the yolk (43.8%) (Fig. 2.C,G). Almost a fifth of them were found deeper in the oocyte (18.8%), among the yolk droplets (Fig. 2.D,H). For the clones whose somatic DNA was not found (n=12/28), we suspect that it was located deeper in the yolk, beyond the first 8 to 10 sections analyzed in our study, from the animal pole downward. Unlike maternal DNA, the injected somatic DNA exhibited different behaviors. In 62 % of the clones, it was organized in metaphasic chromosomes (Fig. 3.B). In 13 % of the clones, a pycnotic pattern was observed (Fig. 3.C). Uncondensed DNA was also present in 25 % of the clones (Fig. 3.D). Interestingly, no clear relationship could be established between the DNA structure and its sub-cellular location in the oocyte (Fig. 3). For example, very well organized metaphasic DNA was unexpectedly observed in the deepest part of the oocyte yolk droplets area, where it should have been less exposed to the MPF-rich ooplasm (Fig. 2.D).

**Figure 3.**
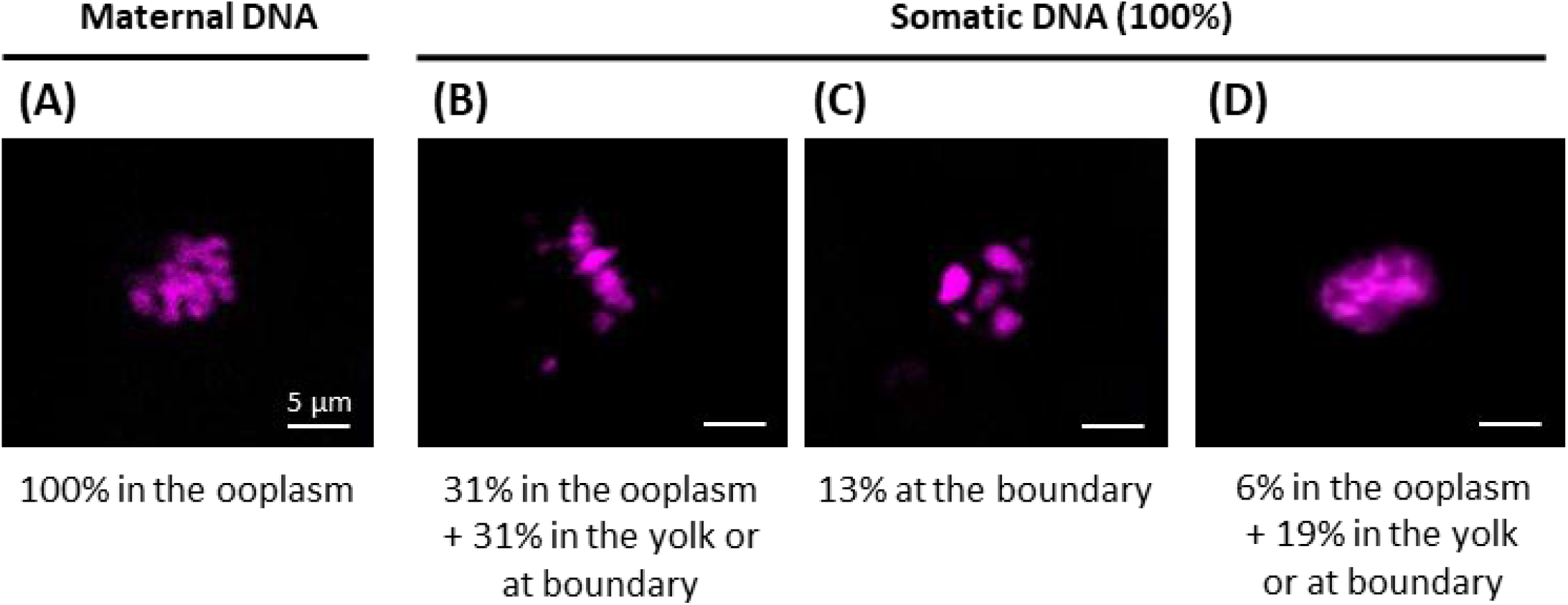
Maternal and somatic DNA structure in non-activated clones, and their respective location within the oocyte. Clones were fixed, cut into 7 μm sections and DNA was stained with Hoechst 33342. Confocal images and localization of (A) maternal DNA condensed into a metaphase plate (MII), (B) somatic DNA condensed into a metaphase plate, (C) pycnotic somatic DNA, and (D) decondensed somatic DNA (interphasic). Those images are representative of 28 clones. Below each image is given the corresponding percentage of DNA found in the ooplasm, at the boundary between the ooplasm and the yolk, and in the yolk.

### Analysis of polar body extrusion pattern after oocyte activation in clones

Meiosis resumption in control oocytes labeled with Hoechst 33342 was characterized by the observation of 3 DNA dots that could be ascribed to the maternal pronucleus, the second polar body and the first polar body (Fig. 4.A). However, neither the shape nor the size of the DNA dots could help us to determine the origin or ploidy of each dot. Therefore removal of the chorion was found to be an efficient way to remove the first polar body as it was no longer attached to the oocyte membrane. Observation of dechorionated samples indeed reduced the DNA dot number to two: the maternal pronucleus and the second polar body. We were able to discriminate between the two dots thanks to the Vybrant Green dye which labeled the second polar body only, while the maternal pronucleus was still labeled with Hoechst (Fig.4.B,C.).

**Figure 4.**
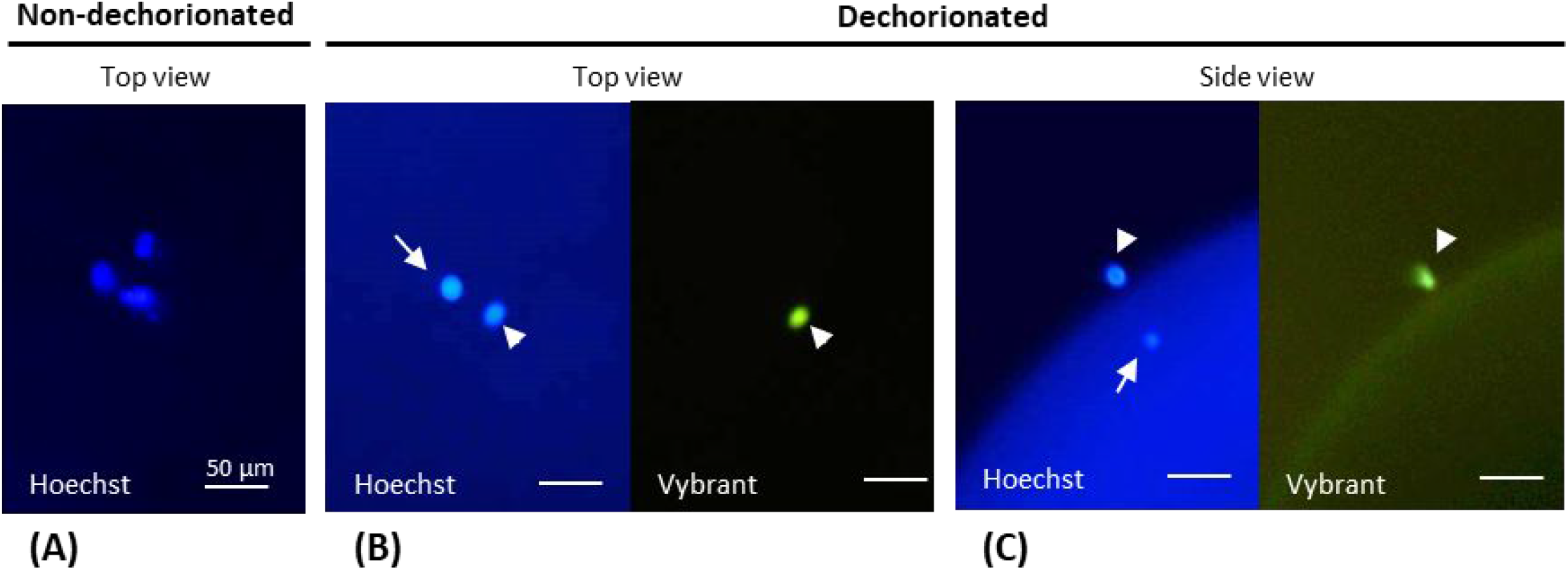
Maternal pronucleus and polar body differential staining in control oocytes after activation. (A) Top view image of a non-dechorionated oocyte: Hoechst 33342 staining did not allow discriminating the maternal pronucleus from the first and second polar body. (B) Top view image of a dechorionated oocyte: the first polar body was eliminated during dechorionation. Vybrant Green dye stained only one of the two DNA dots. (C) Side view image of a dechorionated oocyte: this orientation shows that the second polar body is stained by both dyes, Hoechst 33342 and Vybrant Green, while the maternal pronucleus is only stained by Hoechst 33342. Arrowheads show the second polar body. Arrows show the maternal pronucleus.

After activation, most of the dechorionated control oocytes exhibited only one second polar body, which indicates normal meiosis resumption in untreated oocytes (Fig. 5.A). A similar trend was observed in injected control oocytes, which received the carrier medium but no donor cell. This indicates that perforation of the membrane through the micropyle and injection of the carrier medium did not adversely affect meiosis resumption and polar body extrusion. On the contrary, the rate of extrusion of a single polar body was significantly lower in clones (p=0.001 and p=0.013 in comparisons to control and injected control oocytes, respectively), although it still involved up to 67% of the clones. This result in clones was not correlated with the initial spawn quality as shown by the 24 hours development rate of fertilized controls (Fig. 5.B). Surprisingly, although we have shown that all clones had a normal maternal DNA organized into a regular metaphase plate before activation (see above), almost 20% of them did not extrude any polar body (Fig. 5.A). While all clones underwent a cortical reaction and chorion expansion similar to that of the controls (not shown), such polar body retention mean that SCNT induced a disruption of this specific step of meiosis resumption. Last, we observed some expulsion of two polar bodies in the clones, and surprisingly in the control groups as well (control and injected control oocytes), despite the absence of injected DNA in the latter. In clones, the proportion of two extruded polar bodies was significantly higher than in control oocytes (p=0.002).

**Figure 5.**
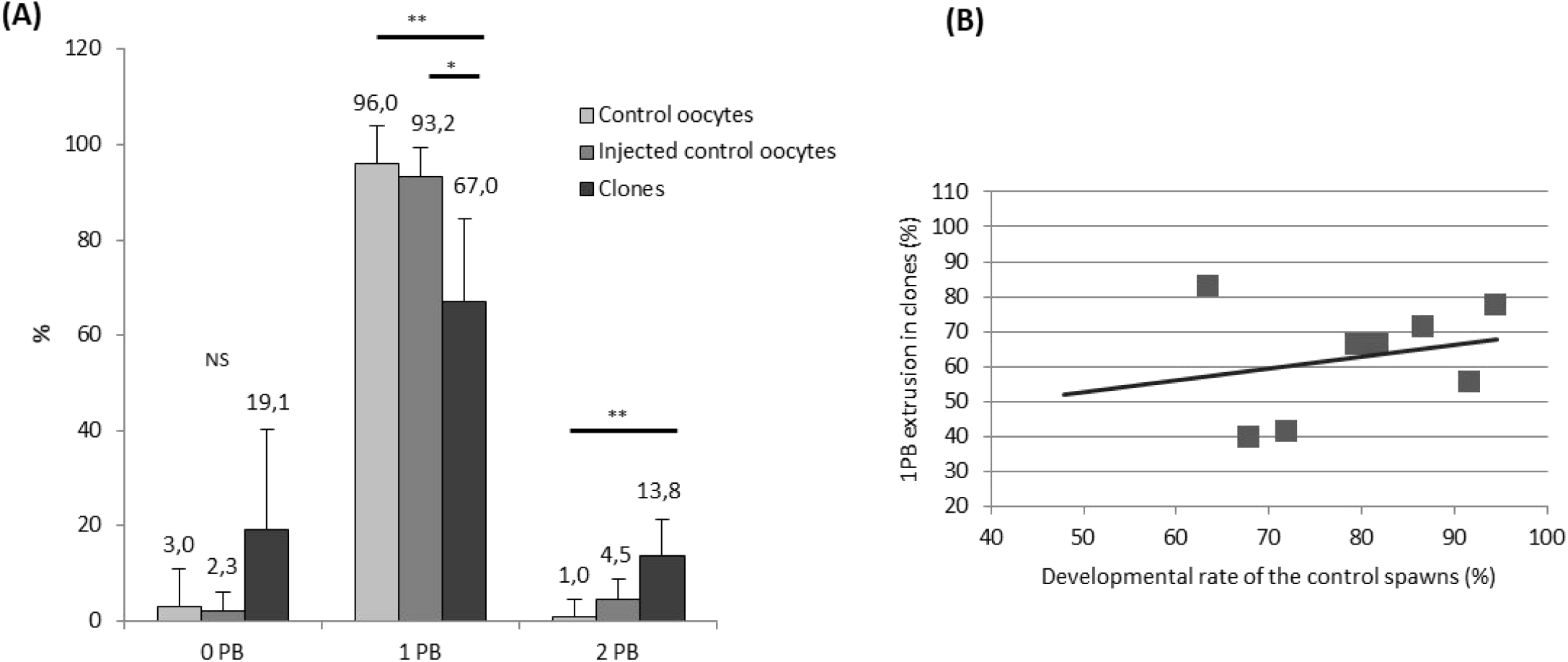
Polar body extrusion after somatic cell nuclear transfer. (A) Graph showing the percentage of activated oocytes which expelled 0, 1 or 2 polar bodies (PB) in control oocytes (from 9 females), injected control oocytes (from 7 females) and clones (from 8 females). Injected control oocytes correspond to oocytes injected with the carrier medium, but without the donor cell. Per females, at least 17 oocytes or clones have been analyzed. NS=Not significant; *=p<0.05; ** =p<0.01. (B) Graph showing the relation between the rate of one polar body extrusion in clones and spawn quality assessed from development rate at 24h post-fertilization. No correlation was observed (r^2^=0.06, n=8 spawns, p>0.05).

### Characterization of DNA fate and blastomeres morphology during the first cell division in clones

The blastomeres morphology during this first cell cycle was characterized in 3 experiments involving a total of 120 clones. Two symmetric blastomeres similar to those of fertilized embryos (Fig. 6.A) were observed in 41 % of clones, while 11 % had 2 cells of asymmetric size (Fig. 6.B). Up to 8 % exhibited 3 or more cells without having gone through an initial 2-cells stage (Fig. 6.C-D). The 3-cell clones always had two symmetric cells and a much smaller one (Fig. 6.C). Up to 40 % of the clones remained in a unicellular state during the observation period (Fig. 6.E). They were not different from the controls that were either activated in water without spermatozoa, or activated in water after injection with the carrier medium without donor cell. However, although those controls collapsed within 2 hours, two thirds of the apparently arrested clones resumed cell division and reached the mid-blastula stage (5 hours post-fertilization, hpf). At the hatching stage (5 days post-fertilization, dpf), 21 % of the clones were still alive. It is interesting to note that almost all these pre-hatched embryos (81%) derived from clones that had shown symmetric blastomeres after the first cleavage.

**Figure 6.**
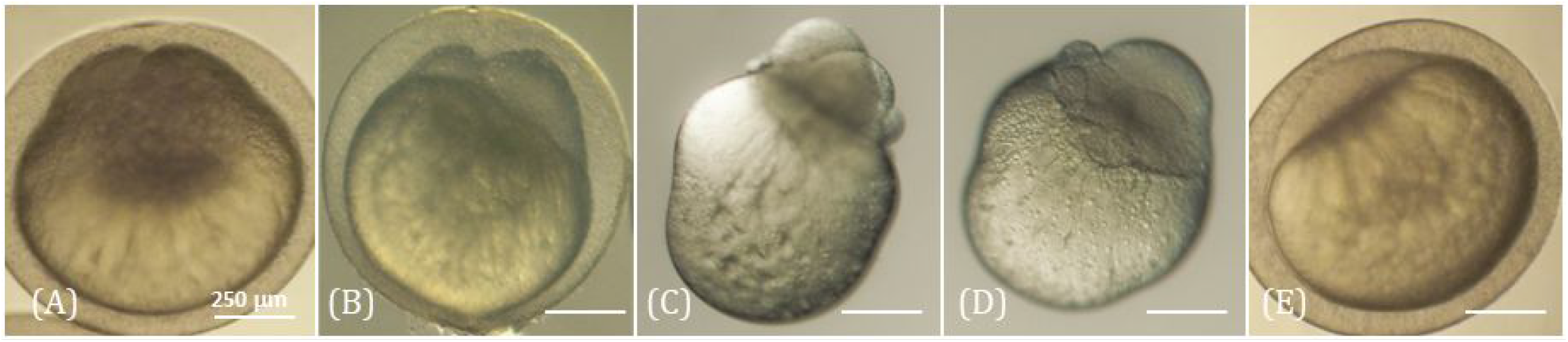
Blastomeres morphology of the clones at the 2 cells stage. Clones were observed 1 h post activation. Images of (A) a non-dechorionated clone with two symmetric blastomeres, (B) a non-dechorionated clone with two asymmetric blastomeres, (C-D) two dechorionated clones with a multicellular configuration, and (E) a non-dechorionated clone which presents no cellular division. Clones were observed under visible light. Morphologies were established on a set of 120 clones.

This better survival of the symmetric clones led us to focus on how the DNA was organized in their blastomeres at the 2 cells stage. We observed in clones and in controls that when the first 2 cells were being formed, the mitosis of the second cell cycle had already begun. As summarized Fig.7.A, despite their correct morphology and cell number, only half the clones with two symmetric blastomeres presented 2 normal mitotic spindles, i.e. a symmetric microtubule spindle bearing DNA in each cell (Fig. 7.B.a,b). Almost a quarter of the symmetric clones presented only one normal spindle in one of the two blastomeres and the remaining clones had only abnormal spindles in their blastomeres. The pattern of these abnormal mitotic spindles was variable: multipolar spindle (Fig. 7.B.c), chromosomal misalignments on the metaphase plate (Fig. 7.B.d), spindle with random DNA location (Fig. 7.B.e), bent spindles (Fig. 7.B.f), and lagging chromosomes resulting from a defect during the first mitosis (Fig. 7.B.g,h).

**Figure 7.**
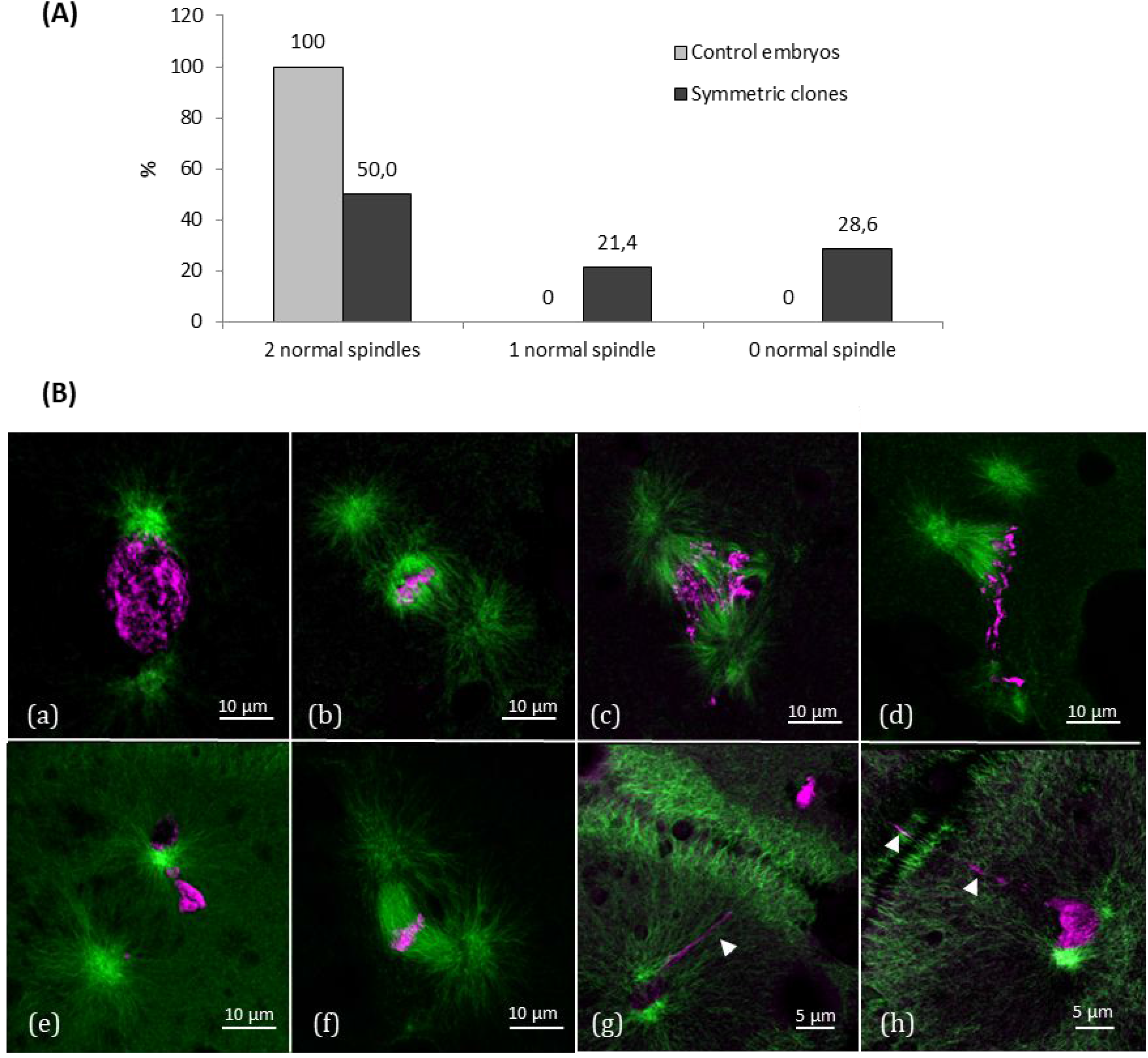
Spindle and DNA structures in symmetric blastomeres of 2 cells stage clones. (A) Graph showing the percentage of clones exhibiting a normal mitotic spindle in both blastomeres (2 normal spindles), only one normal spindle in one of the two cells (1 normal spindle), or only abnormal spindles (0 normal spindle) (n=7 control embryos, n= 14 symmetric clones). (B) Confocal images of normal and abnormal spindles in clones. Embryos were fixed, and cut into 7 μm sections. Microtubules were immunolabeled with an anti-α-tubulin antibody (green) and DNA was stained with Hoechst 33342 (magenta). Images of (a) a normal figure with duplicated centrosome at each side of the prometaphasic DNA, (b) a normal spindle with condensed metaphasic DNA, (c) an abnormal tripolar spindle with anaphasic DNA, (d) an abnormal multipolar spindle with metaphasic DNA, (e) an abnormal spindle with unidentified DNA stage and localization, (f) an abnormally bent spindle with metaphasic DNA, (g-h) lagging chromosomes (arrow heads) spanning accross the cleavage furrow of the first mitosis.

Strikingly, almost two-thirds of the symmetric clones (11/14; 78%) showed fragmented DNA under the cleavage furrow separating the first two blastomeres (Fig. 8), while no such figure was observed in control embryos (0/7). This fragmented DNA presented no clear condensed chromosomes, nor centrosomes or a microtubules network.

**Figure 8.**
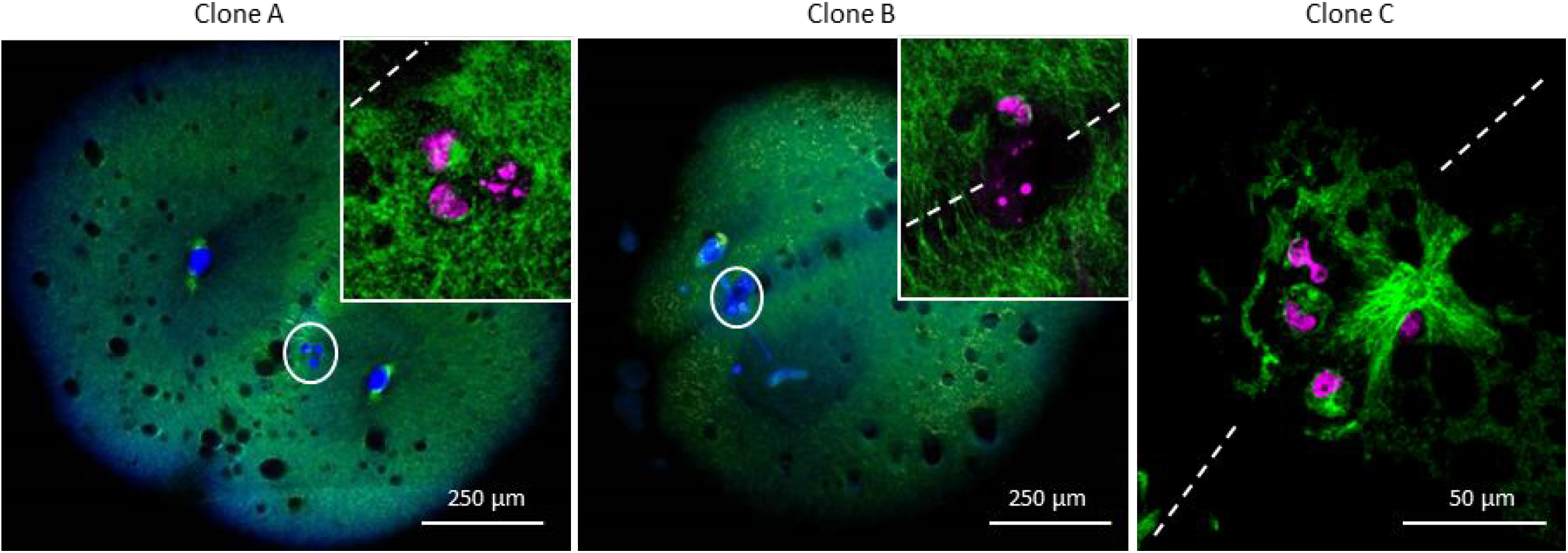
Identification of fragmented DNA on the cleavage furrow of the clones. Embryos were fixed, and cut into 7 μm sections. Microtubules were immunolabeled with an anti-α-tubulin antibody (green) and DNA was stained with Hoechst 33342 (blue or magenta). Reconstitution of several section images of two clones at 2-cells stage (Clone A and B). White circles: fragmented DNA (blue) located on the cleavage furrow. Insets and clone C: images of the fragmented DNA (magenta) observed in by confocal microscopy, dotted lines show the cleavage furrow. Those images are representative of 11 clones.

## Discussion

The question of chromatin fate after SCNT has always been a matter of acute concern in all studied species, although the complexity of most fish eggs hindered the access to such knowledge to date. In our study, we provide the first information on the behavior of the maternal and somatic chromatins in a system where the oocyte (maternal) DNA was not removed prior to SCNT.

### Fate of the maternal DNA after SCNT

This study showed that prior to activation of embryonic development, the maternal DNA at MII stage remained undisturbed by the injection procedure of SCNT. Its location against the micropylar canal probably protected the maternal DNA from the injection needle and its integrity was not compromised by nuclear transfer. Finding this chromatin against the canal and not at its bottom, at the sperm entry site, may also explain why successful enucleation by aspiration at this oocyte stage has never been reported in fish. This sheltered position also rules out the hypothesis that the maternal DNA could be stripped off or damaged during the somatic cell injection through the micropyle. Our results indeed demonstrated that maternal DNA loss did not occur at this step of the process. Moreover, this homogeneity of maternal DNA structure and location in clones cannot account for the variability in the embryo fate observed later on during development.

Our finding that the maternal DNA metaphase was undisturbed by SCNT was making it likely that extrusion of a second polar body of maternal origin would be maintained in our system. As thoroughly studied in mouse, polar body extrusion requires several conditions including chromosome condensation in a metaphasic plate and its location close to the plasma membrane where interaction with the cortical cytoskeleton will trigger asymmetric cleavage^24,25^. And indeed, polar body extrusion was maintained in most clones, so we infer that the maternal DNA was always involved. Therefore, the microcapillary penetration or the TCF carrier medium injection did not alter substantially the integrity of the cortical cytoskeleton that is necessary for the maternal polar body formation. The extrusion of an additional polar body (two extruded polar bodies) that was observed in some cases can be ascribed to the somatic DNA, as will be discussed later. It was surprising however that a small part of the control oocytes would also extrude two polar bodies, although no somatic DNA was present. This phenomenon could be explained by an incorrect positioning of the meiotic spindle at the plasma membrane upon meiosis resumption, leading to the total expulsion of maternal DNA in two distinct globules, as already described in mouse after the microtubules network had been deliberately destabilized^24^. If this defect was to occur in clones as well, it may be responsible for a small number of embryos undergoing spontaneous loss of maternal DNA upon meiosis resumption. To date, we did not identify the genetic origin of the polar bodies to develop further this hypothesis. Last, the rate of clones with no polar body extrusion was higher than in control groups, although this was not statistically significant because of high variability between cloning series. It means that in some series, the somatic cell injection was deleterious and prevented the polar body extrusion process. The hypothesis of an intracellular signaling defect, such as a disturbance of the intracellular concentration of calcium, could be a reason for polar body extrusion failure in some cases as reported previously in mouse^26^. Since we injected a whole somatic cell, its own cytoplasmic cargo was injected as well, which could have disrupted the activation signaling pathway, but we have no clue about why this alteration would occur in some cloning series and not in others, as this observation was not related to spawn quality.

### Fate of the somatic DNA after SCNT

Contrarily to the stability demonstrated for maternal DNA before oocyte activation, the structure of somatic DNA was more variable, although it still adopted a metaphasic pattern in two third of the clones. It has been previously shown that even though the whole cell is injected, its plasma membrane is disrupted within the first few seconds following SCNT^22^. Therefore, the somatic DNA is readily exposed to the high MPF levels of the oocyte. It was therefore expected that the chromatin would adopt a metaphasic stage, despite the fact that the injected nuclei in G0/G1 phase had not undergone its own S phase (DNA synthesis) and that the somatic DNA had not been duplicated. Such condensation pattern upon injection into a non-activated oocyte resembles the so called Premature Chromatin Condensation (PCC) phenomenon described in mammals, whose favorable or unfavorable somatic reprogramming efficiency and contribution to the success of embryonic development is still a matter of debate^27–33^. But the similarity ends up there, because in mammals, extrusion of a second polar body that would halve clone ploidy is prevented by cytochalasin B^34,35^. This procedure has never been used in fish, as penetration and release of chemicals in the highly amphiphilic fish eggs is difficult to achieve^36^, and cytochalasin B trapped in the egg is deleterious for ooplasmic segregation at the onset of embryonic development^37^. At this point, two scenarios can be proposed in fish clones: for those clones whose condensed somatic DNA was close enough to the oocyte plasma membrane prior to meiosis resumption, their chromosomes may have been able to interact with the cortical cytoskeleton, resulting in the extrusion of an additional polar body from somatic origin. We reported that 31 % of the clones had a condensed somatic DNA localized into the ooplasm. Some of them could count among those which expelled two polar bodies, supposedly from both DNA origins. In this case, the clones would end up with a haploid somatic DNA and a haploid maternal DNA, resulting in a diploid hybrid. This was not assessed in the present study, but such diploid hybrids were reported in medaka^18^ and in goldfish (Labbé et al, personal communication). The second scenario is when the somatic DNA was too far from the plasma membrane to be able to contribute to polar body extrusion. The odds for this scenario are favorable in fish when considering the huge size of the oocyte and the low surface/volume ratio at the animal pole. This is strikingly supported by a study on mouse oocyte^28^ in which injected DNA beads underwent ectopic polar body extrusion only when they were close enough to the cortex, showing a distance-dependent induction of polar body extrusion. At this point, the fate of the somatic DNA upon activation and meiosis resumption signaling is difficult to ascertain from our data. Somatic chromosomes could undergo a pseudo-anaphase that would lead to the formation of two haploid nuclei in the clones, or the pseudo-mitosis would not operate on the somatic nucleus, despite the loss in MPF, and a single diploid nucleus would reform thereafter. We believe that these options are the roots for the high variability in mitotic figures during early development.

But we also reported that not all clones bore somatic DNA in a metaphase stage, as almost one fourth of them was found uncondensed in the oocyte. We infer that these clones would skip the meiosis resumption process, and thereafter resume the embryonic cell cycle leading to regular DNA replication and first mitosis. Those clones may account for the successful diploid clones reported in many studies^18–20^. However, we could wonder if the clones whose somatic DNA location is deeper in the oocyte will be able to develop. Indeed, the somatic DNA was sometimes observed among the yolk droplets. If this DNA was to contribute to the embryo development, it would have to migrate towards the animal pole, possibly within the powerful cytoplasmic streams observed^38^ during the first cell formation. Although not precisely assessed here, this hypothesis may be one explanation for those clones showing one seemingly stalled blastodisc at the 2 cells stage but which developed later on and reached at least the mid-blastula stage. In these, the somatic DNA would be delayed at reaching the ooplasm where embryonic mitosis could finally resume.

### Fate of the maternal DNA upon first cell cleavage

From the above discussion, several clones should still possess the entire diploid somatic DNA, either because it was uncondensed, or because it was condensed but far from the plasma membrane. And because of the successful maternal meiosis resumption, a high majority of the clones should still possess half the maternal DNA as well. This raises the question of how this maternal DNA behaved during clone development, especially during the first mitosis, and whether it interfered with the somatic DNA. It has been proposed in medaka that the presence of maternal DNA is beneficial for the cellular reprogramming of the somatic cell, by decreasing cleavage asynchronies and ploidy mosaicism in clones^19,39^. However, no study provided any hint about how this would operate in the first embryonic cells.

One striking observation in the present study is that most symmetric clones displayed a fragmented DNA that was located under the cleavage furrow of the first cell division. It is known that the position of the cleavage furrow is determined by the positioning of the mitotic apparatus, the latter being accurately centered in the fish egg blastodisc by dynein-associated pulling forces^40,41^. Polar body extrusion also takes place in a centered position, at the apex of the animal pole^40,42^. It means that in our system, the cleavage furrow is a landmark of where half the maternal DNA was extruded and half remained in the embryo. The fact that some DNA was found at the bottom of the cleavage groove strongly suggests that it is of maternal origin and that it remained at its original location in the absence of any capture by some microtubular apparatus before the first embryonic cleavage. In mutant zebrafish where male centrosome was made unable to attach the maternal pronucleus, the later did not undergo any migration either, and it stayed at its original location close to the polar body^43^. In other mutants, ectopic masses of DNA at early stage, that can be compared to the maternal DNA in our clones, subsequently became fragmented or got lost^44^. In all, this could explain how in some fish clones, the maternal DNA was sequestered and eliminated during subsequent mitosis.

### Fate of the clones during embryonic development

Even if DNA of maternal origin was excluded during development in some clones, embryonic development still stayed vulnerable when considering the wide set of alterations reported here during the first cell cycle. Symmetry defects could be due to a miscentering of the first mitotic spindle, because of altered interactions of the somatic mitotic spindle with the microfilament network of the blastodisc in the clones. But even when clones successfully underwent a first symmetric cleavage, we report that spindle defects and chromosome lagging and misalignments persisted in many of them. Because an entire somatic cell was injected, its own centrosome (centrioles and pericentriolar material) and its own cytoplasmic cargo (proteins and RNA) were incorporated and could have induced a protein or RNA imbalance of the oocyte cargo. One nagging question remains about the interaction between the somatic centrosomal apparatus and maternal pericentriolar material. The broad time window (> 40 min) of this event taking place before the first cleavage impeded our ability to explore it in the present study. We believe nevertheless that the injected somatic centrosome may have contributed to successfully replace the missing spermatic one.

### Conclusion

To summarize (Tab. 1), we demonstrated that the maternal metaphasic DNA was undisturbed after SNCT and that microinjection of a somatic cell through the micropyle was not impeding the oocyte plasma membrane ability to support polar body extrusion. Moreover, we provide strong indications that during mitosis, the maternal DNA remained at its original position, resulting in its positioning under the first cleavage furrow of the clones and its likely dispersion later on at the interface between yolk and embryonic cells. We also reported a high variability in somatic DNA structure and location which may account for the high variability in clone ploidy and blastomeres morphology during development. We also proposed some hypothesis about how clone ploidy could be maintained in some cases in relation to the somatic DNA structure and location upon injection, and despite the fact that no cytochalasin B is used in fish SCNT. This is the first time that some information on the cellular events taking place after SCNT is provided in such a specific model that is fish oocyte and embryo.

**Table 1.** Summary of the results regarding the fate of the maternal DNA prior oocyte activation, after meiosis resumption and after the first mitosis. (MII : oocyte metaphase 2). +++: > 90 %; ++: > 50 %; +: > 10 %; +/-: > 0 %; *: mostly (81 %) from symmetric cleavage.

|  | CONTROLS | CLONES |
| --- | --- | --- |
| Maintenance of maternal MII | +++ | +++ |
| **MEIOSIS RESUMPTION** |  |  |
| 1 PB extrusion | +++ | ++ |
| 2 PB extrusion | +/- | + |
| **MITOSIS** |  |  |
| Symmetric cleavage | +++ | + |
| Survival at hatching | +++ | +* |

## Materials and methods

### Gametes collection

Two years old mature goldfish (*Carassius auratus*), originating from outdoor ponds at INRA U3E experimental facility (Rennes, France), were used as breeders. Male and female were kept in separated 1m^2^ tanks, at a constant temperature of 14°C, under spring photoperiod (16h light and 8h dark cycle). They were transferred at 20°C few days prior to hormonal induction by intra-peritoneal injection of 0.5 mL/kg Ovaprim™ (Syndel Laboratories, Canada). Sixteen hours later, gametes were collected by stripping. Oocytes were maintained at 12°C for up to 5 hours in their own coelomic fluid. Sperm was diluted in SFMM (NaCl 110 mM, KCl 28.3 mM, MgSO_4_ 2H_2_O 1.1 mM, CaCl_2_ 2H_2_O 1.8 mM, Bicine 10 mM, Na Hepes 10 mM, pH 7.8, 290mOsm/kg) and stored on ice for up to 24h. Fish handling and sampling was carried out in strict accordance with the welfare guiding principles of the French regulation on laboratory animals, under the French and European regulations on animal welfare (Authorization N° 005239 level 1; C. Labbé), and under the supervision of staff of the animal facilities possessing an agreement level (C35-238-6). The experimental protocol was approved by the welfare committee from the Fish Physiology and Genomics department at Institut National de la Recherche Agronomique (registration C-2018-01-CL-AD) in accordance with the French guidelines on broodstock handling, gamete collection and embryo rearing.

### Nuclear transfer procedure

Nuclear transfer was carried out as described previously^22^. Donor somatic cells were obtained from caudal fin after explant culture and cell cryopreservation^14^. After thawing, mesenchymal cells in G0/G1 stage were washed in the cell culture medium with antibiotics (2.5 μg/mL amphotericin B, 50μg/mL gentamicin) and stored on ice for up to 2 hours. Nuclear transfer was performed at 20°C with a Cell Tram Vario injector (Eppendorf) connected to a micromanipulator (Transferman NK2, Eppendorf) under a stereomicroscope (Olympus SZX 12). Recipient oocytes were placed into Trout Coelomic Fluid (TCF) to prevent oocyte activation^45^. Donor cells were isolated in TCF as well. After localization of micropyle at the animal pole (sperm entry point), oocytes were held by gentle depression of the holding microcapillary (iD 100 μm). A single donor cell was aspirated in a glass microcapillary (iD 15 μm, custom Tip Type IV, Eppendorf) and injected with about 10 pL of TCF into the recipient oocyte through the micropyle, just under the oocyte plasma membrane.

After nuclear transfer, oocytes were incubated for 30 min in TCF to improve success rate of the nuclear transfer^22^. They were then either fixed for analysis, or activated with tap water to trigger embryo development. Embryos were incubated at 20°C in dechlorinated tap water. When specified, the chorion was enzymatically removed by incubation of the embryos with 4 mg/mL protease from *Streptomyces griseus* (SIGMA, P8811) diluted in Holfreter 2.2 (NaCl 60 mM, CaCl_2_ 2H_2_O 68 μM, KCl 67 μM, D-Glucose 12 mM, PVP 40 000 62.5 μM, Hepes 5 mM, pH 7.4, 140 mOsm/kg). Dechorionated embryos were incubated in Holfreter 2.2 at 20°C up to fixation and analysis. In analyzes devoted to the clone fate, several development stages were specifically observed: the 2 cells stage (about 1 hpf), when a scission grove appeared on the top of the single blastodisc, the mid-blastula stage (5 hpf) when the control embryos reached the 1000 cells stage and began embryonic layers differentiation, and the hatching stage (4-5 days post fertilization, dpf), when the control embryos were released from the chorion.

At the end of each SCNT session, oocyte quality was checked by an in vitro fertilization test. About 100 oocytes from each of the spawn used for nuclear transfer were fertilized in dechlorinated tap water with 10 μL diluted sperm. The number of live embryos was assessed at 24 hpf, 6-9 somite stage and at hatching, and expressed as a percentage of the initial oocyte number. Oocytes were graded as good quality when the development rate was above 90 % at 24 hpf. Nevertheless, all experiments whose control rates were below 60% were discarded.

### Fixation and immunofluorescence analysis after nuclear transfer

After nuclear transfer, non-activated oocytes and embryos (2 cells stage) had to be orientated with the animal pole up, to ensure identification of the DNA on sections during histological analysis. Oocytes were orientated prior to methanol fixation, when the micropyle used to identify the animal pole was still visible. To prevent activation, oocytes were held in 2 % gelose (agar-agar, PROLABO 20768.235) cupules prepared in Goldfish Ringer medium (GFR: NaCl 125 mM, CaCl_2_ 2H_2_O 2.4 mM, KCl 2.4 mM, MgSO_4_ 7H_2_O 0.3 mM, MgCl_2_ 6H_2_O 0.9 mM, D-glucose 6 mM, Hepes 4 mM, pH 7.3, 256 mOsm/kg) supplemented with Soybean Trypsin Inhibitor Type II-S (SIGMA, P9128), and orientated under binocular with the micropyle up. They were then covered with melted 2 % agar-agar gelose, cooled, and fixed with cold methanol overnight.

Dechorionated embryos at 2 cells stages were directly fixed with cold methanol overnight and all samples were stored at −20°C until gelose embedding. Methanol prevented blastodisc and blastomeres deformation and was mandatory for immunofluorescence specificity. After fixation, early embryos were transferred in 70 % ethanol to limit methanol toxicity, orientated with the embryonic cells up, and embedded in a 2% agar-agar gelose prepared in distilled water.

For paraffin embedding, oocytes and embryos in their agar cushion were transferred in 100 % ethanol, 96 % ethanol (2 x 30 min), butanol-1 (3 x 3 h and 1 x 1 h) and melted paraffin (60°C, 2 x 2 h). Samples were then embedded with the animal pole up in paraffin cassettes and cut into 7 μm sections. Only one fourth of the sample was cut for histological analysis (approximatively 250μm). This area corresponds to the ooplasm. Sections were then mounted on slides with ovalbumin 0.5 % (PROLABO, 20771.236). Paraffin removal and rehydration of the slides was carried on in successive bath of toluene 100 %, ethanol (100 %, 96 % and 70 %), and PBS (SIGMA, P4417).

For fluorescence immunolabeling, all slides were saturated in PBS with 2 % bovine serum albumin (BSA, SIGMA, A2153) and 0.5 % Triton X-100 (SIGMA, T8787)(1.5 h, 20°C) and incubated with mouse anti-tubulin-α antibody (SIGMA, T9026)(1.5h, 30°C). After washing in PBS-BSA 0.2 %, samples were incubated with the goat anti-mouse antibody coupled with Alexa-Fluor 488 (SIGMA, A11001) (1.5 h at 30 °C). After washing, samples were stained with Hoechst 33342 (SIGMA, B2261) (2.5 μg/mL, 15 min, 20°C). Slides were mounted with PBS-Glycerol solution and stored at 4°C until analysis. Fluorescence observation and images were taken under a fluorescent microscope (Nikon 90i) and a SP8 confocal microscope (Leica microsystems) to characterize the DNA structure and localization.

### *In vivo* study of polar body extrusion

In order to visualize the polar body extrusion, oocytes were stained with two live DNA markers: Hoechst 33342 labeled both the maternal pronucleus and the extruded polar body, while Vybrant Green dye (Invitrogen, V35004) labeled only the extruded polar body. We believe that the amphiphilic Vybrant dye penetrated the oocyte well, but the composition of the ooplasm prevented this dye from specifically bind the maternal DNA. Instead, the entire ooplasm displayed a very weakly non-specific green fluorescence. This bias was used for our purposes because only the DNA incorporated into the tiny cell that is the polar body could be labeled with Vybrant Green, while the maternal pronucleus was only labeled with Hoechst. Non-activated oocytes were incubated with 100 μg/mL of Hoechst 33342 in TCF for 30 min at 20°C, and rinsed twice with TCF to prevent non-specific Hoechst signal. After oocyte activation in dechlorinated tap water for 1 min, chorions were removed as described above within 10 min to ensure polar body visualization. Dechorionated oocytes were incubated with 10 μM Vybrant green dye in Holfreter 2.2 medium, before and during the polar body extrusion (around 13 min post-activation at 20°C). To prevent the loss of the polar body, oocytes were no longer manipulated from this time on. Observation of the polar body could be carried on under a stereomicroscope up to 40min post-activation.

### Statistical analysis

Data were expressed as mean ± standard deviation. One-way ANOVA followed by Kruskal-Wallis test (non-parametric test) was performed using IBM SPSS Statistics, Version 24.0. Armonk, NY:IBM Corps.

## Acknowledgements

The authors thank the staff from INRA U3E (Rennes) for providing goldfish breeders. F. Borel, A. Patinote, J-M. Aubry, C. Duret and P-L. Sudan took great care of the goldfish at the INRA LPGP experimental facility (Rennes). The members of the INRA LPGP histology service, A. Branthonne and B. Porcon are also acknowledged for training and technical support. G. Halet, from the Institute of Genetics and Development (IGDR Rennes), contributed to the reflection and results deciphering thanks to his fundamental knowledge of the meiotic and mitotic divisions. N. Beaujean (INRA-INSERM SBRI, Lyon) also helped us to deepen our interpretations thanks to her knowledge on cellular events after nuclear transfer. The other members of the thesis committee are also thanked for their kind help and support. Audrey Laurent (INRA LPGP Rennes) provided valuable comments on the manuscript and interpretation of the data. This work has benefited from the Tefor Fish Phenotyping Platform at the INRA LPGP, Rennes (ANR-II-INBS-0014) for confocal analysis where the expertise of V. Thermes and M. Thomas is gratefully acknowledged. This work was funded by the French CRB Anim project, ANR-11-INBS-0003. CR was recipient of an INRA PHASE and Région Bretagne PhD fellowship.

## Author Contributions

CR organized and carried out the study, analyzed and interpreted the data and drafted the manuscript. AD performed the nuclear transfer experiments and contributed to the coordination of the study. NC provided the donor somatic cell culture, initiated the immunofluorescence protocols and participated to the clone fate experiment. PYLB and CL conceived and designed the study. CL supervised the experiments and the manuscript writing. All authors read, improved and approved the final manuscript.

## Additional Information (including a Competing Interests Statement)

The authors declare no competing interests.

